# Live cell bioimaging with carbon dots produced in situ by femtosecond laser from intracellular material

**DOI:** 10.1101/455808

**Authors:** Aleksander M. Shakhov, Artyom A. Astafiev, Alina A. Osychenko, Maria S. Syrchina, Viktor A. Nadtochenko

## Abstract

Owning to excellent optical properties and high bio-compatibility carbon dots (CDs) have drawn increasing attention and have been widely applied as imaging agents for various bio-applications. Here we report a strategy for live-cell fluorescent bioimaging based on in situ synthesis of CDs within cells by tightly focused femtosecond laser pulses. Laser-produced carbon dots exhibit bright excitation-dependent fluorescence and are highly two-photon active under near infrared femtosecond excitation, thus demonstrating a potential for two-photon fluorescence imaging. The Raman spectra of fluorescent centers show strong D (1350 cm^−1^) and G (1590 cm^−1^) bands, thus suggesting that they are composed of carbon dots with sp^2^-hybridyzed core. Using Mouse GV oocytes as a model system we examine cytotoxicity and demonstrate the possibility of long-term fluorescent intracellular tracking of the laser-produced CDs. Created virtually in any point of the live cell, CD-based fluorescent µm-sized markers demonstrate high structural stability and retain bright fluorescence many hours after formation. Our results point to laser-produced fluorescent CDs as a highly-potent tool for cell cycle tracking, culture cell marking and probing intracellular movements.

## INTRODUCTION

Cell cycle and intracellular movement visualization are of much interest in biomechanics and biochemistry. Fluorescent bioimaging is widely used to visualize cell structure and dynamics. Usually fluorescent imaging relies on fluorescent molecules – organic dyes or fluorescent proteins. However, molecular fluorophores suffer from a number of limitations: they are subject to photobleaching and influence of microenvironment on their fluorescent properties, typically have a small Stokes shift and are frequently phototoxic. For this reason, development of fluorescent nanomaterials, which could be used as an alternative to organic dyes and fluorescent proteins in bioimaging, has attracted much interest. Doped nanoparticles, semiconductor quantum dots, metal nanoparticles, polymer dots, carbon quantum dots, dendrimers, etc. were proposed as promising nanomaterials [1] that offer advantages of greater photostability, lower toxicity, inertness in the intracellular environment, and more advantageous spectral properties compared with molecular fluorophores [2]. Among various fluorescent nanomaterials carbon dots or carbon nanodots possess an optimal combination of bright visible fluorescence [3], photostability [4], excellent biocompatibility [5] and inexpensive and environment-friendly production technology, which makes them a perfect choice for bioimaging applications [6].

A number of methods for carbon dots (CDs) production have been proposed which can be broadly divided into top-down cutting of carbon materials [7-10] and bottom-up synthesis [11-13] from a variety of organic precursors including even food waste [14]. Virtually all known methods demonstrate production of relatively large quantities of material in a macroscopic volume. Also the methods of intracellular bioimaging with fluorescent CDs rely on internalization of exogenous particles from external media. Unlike organic dyes, nanoparticles are not capable of permeating cell membranes, and their internalization occurs mainly through various mechanisms of endocytosis [15-17]. These mechanisms are still not fully understood and often have a limited effectiveness, which also varies greatly depending on characteristics of the nanoparticles, the type and state of the cell, and the type of the medium. Control of dynamics and intracellular localization of nanoparticles presents an additional problem from a standpoint of their targeted delivery [18-19]. Synthesis of carbon dots directly within cells from intracellular organic molecules, while possible in principle, has not been demonstrated thus far.

Controlled modification of material properties in a microscopic volume can be realized by means of nonlinear absorption of ultrafast laser pulses. This approach was widely applied in technologies of laser micro- and nanoprocessing of materials [20-22], including laser-induced chemical modification such as laser polymerization [23-24] or multiphoton photoreduction of metal ions [25-26]. As concerns biological materials, nonlinear optical interaction with ultrafast laser pulses were studied mostly from a perspective of laser micro and nanosurgery of cells and tissues [27-28]. It was demonstrated that mechanisms of laser surgery include chemical reactions induced by electronic plasma generated by nonlinear photoinization. In principle, similar to a plasma formed by electric discharge [29-30], laser-generated microscale plasma can stimulate carbonization of organic materials and formation of luminescent carbon dots. It was reported that irradiation with pulsed laser light can induce synthesis of CDs from an aromatic precursor possibly through a plasma-mediated mechanism [31]. While there were evidences of femtosecond laser-induced carbonization in biological tissues [32-34] and formation of light absorbing centers [35] a laser synthesis of carbon dots from bio-molecules has not been positively demonstrated.

In this paper we report that ultrafast laser processing can be employed to generate fluorescent carbon dots in intracellular material through nonlinear absorption of tightly focused femtosecond laser pulses. These carbon dots are localized in a microscopic volume down to a submicrometer size, and exhibit high chemical stability and low cytotoxicity. We demonstrate that laser-generated CDs-based microscopic fluorescent markers can be employed to trace material reallocation during oocyte maturation. Our findings highlight viability of bottom-up synthesis of carbon dots from organic precursors in high-intensity optical field of laser pulses and its application for fluorescent biological imaging.

## MATERIALS & METHODS

Detailed description of experimental procedure is given in the Supporting Information. In short, NIR femtosecond laser pulses at 80 MHz repetition rate from a Ti:Sapphire oscillator were coupled to an optical microscope and focused by an objective lens (0.75 NA) into a focal spot of 1.3 μm diameter. Mouse GV oocytes in PBS solution inside a Petri dish were placed on a microscope sample stage and were irradiated with focused femtosecond pulse trains with a length of 20-200 ms and a pulse energy up to 3 nJ (peak intensity up to 9 MW/cm2). Optical image of the oocytes was recorded with a video camera, for fluorescent imaging oocytes were illuminated with a CW laser diode at 462 nm, and excitation light was blocked by a LP500 spectral filter. Fluorescence emission spectra and decay kinetics of intracellular material were registered locally in the focal spot area using two-photon excitation with femtosecond laser light at low power. Oocyte material’s Raman spectra were excited with a 532 nm CW laser (DPSS, Coherent) and recorded with a Raman spectrometer Renishaw 1000B attached to the microscope. After femtosecond laser irradiation oocytes were kept in a CO2 incubator for a period up to 24 hours and were inspected with the optical microscope thereafter.

## RESULTS & DISCUSSION

Irradiation of the oocyte material with a train of tightly focused femtosecond laser pulses resulted in changes in material’s fluorescence. Normally the intracellular material is only weakly fluorescent due to presence of endogenous fluorophores. However, after laser irradiation a well-visible brightly fluorescent micrometer-sized spot was formed around the laser focal point (Fig. 1a, b). Fluorescence intensity in the bright spot was typically many folds higher than in non-irradiated material. This effect indicates that considerable quantities of fluorescent products are generated in the intracellular material under femtosecond laser irradiation. Accumulation of fluorescent products was also monitored with fluorescence excited by two-photon absorption of femtosecond light. Figure 1c presents a dependence of fluorescence intensity recorded during a 135-ms exposure to the laser pulses. Whereas the initial intensity was negligible, there was a many-fold nearly monotonous increase in emission which reached the maximal value at the end of exposure. This effect indicates that the generation of fluorescent material is gradual and continues during the entire period of laser exposure.

**Figure 1.**
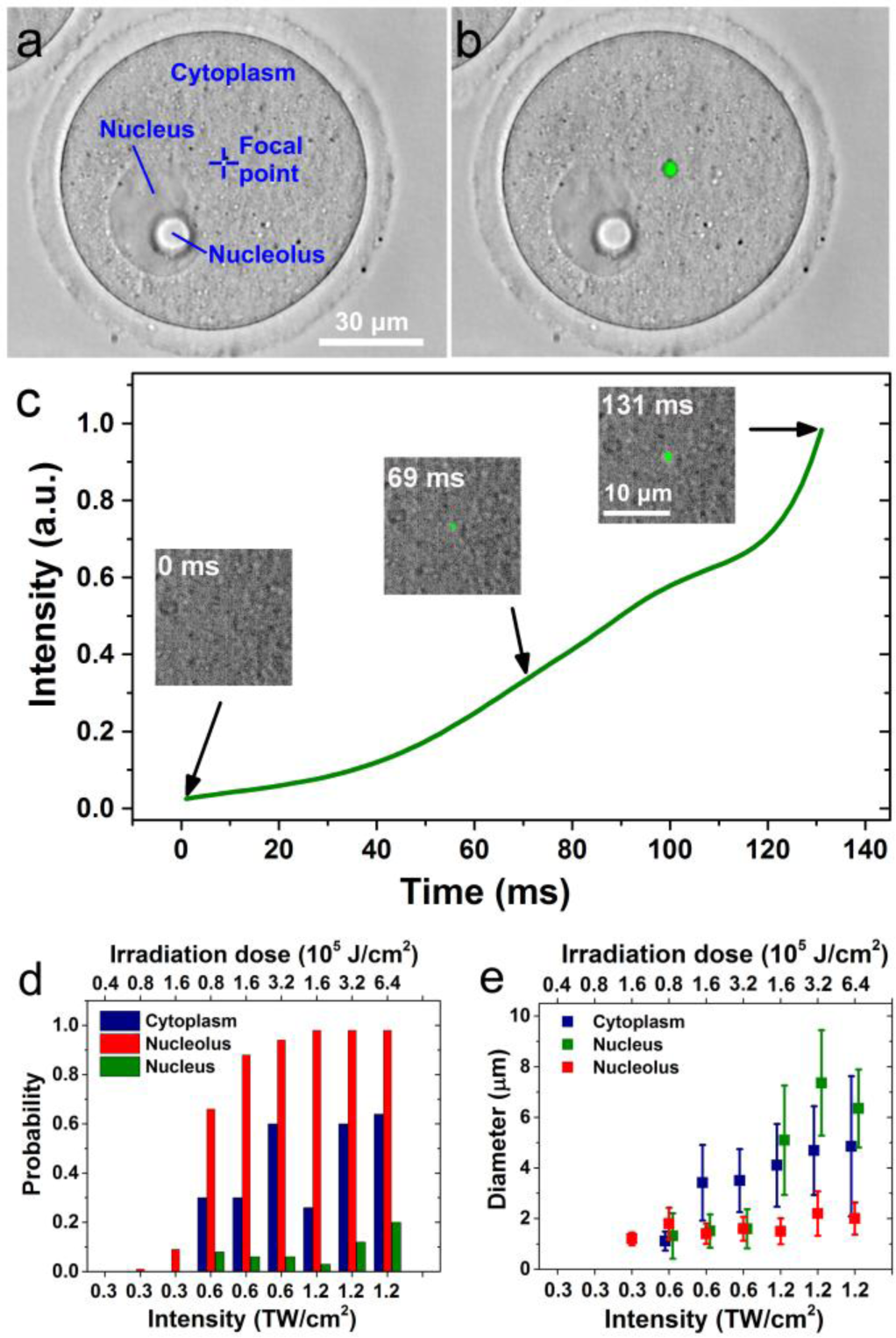
Images of the oocyte (**a**) before, and (**b**) after exposure to a laser pulse train, combination of brightfield and fluorescent image, fluorescence is shown in green color. Fluorescence was excited by a laser diode at 462 nm. Laser intensity was 1.2 TW/cm^2^, train length – 100 ms. Characteristic diameter of the fluorescent spot − 4 μm. (**c**) Intensity of two-photon excited fluorescence from the irradiated area during exposure to a train of pulses at 1.2 TW/cm^2^ intensity vs. time. Insets show respective images of the oocyte material around the laser-irradiated spot, green color shows laser-induced fluorescence. (**d**) Probability of fluorescent spot formation as a function of irradiation parameters (**e**) Average diameter of the fluorescent spot as a function of irradiation parameters, absence of points means that fluorescent spot formation was not registered.

Similar effects of fluorescent products generation were observed in three distinct regions of the oocytes with different chemical composition: cytoplasm, nucleus and nucleolus. Formation of fluorescent spots had a threshold-like dependence on laser pulse parameters: it required sufficiently large peak laser intensities and irradiation doses (Fig. 1d). For example, there were virtually no effects of fluorescent material generation at 0.3 TW/cm^2^ intensity, whereas these effects were visible at larger intensities. The characteristic diameter of the fluorescent spot varied greatly in different points of the oocyte even at the same irradiation parameters probably due to variation in absorption. Nevertheless, we observed a tendency for spots to become larger with an increase in laser intensity and irradiation dose (Fig. 1e). With intensity and dose close to the formation threshold we could produce fluorescent spot of sub-micrometer diameter which was comparable to the laser beam focal spot size (Fig. 1e and Fig. S1).

Fluorescent products generated in cellular material by femtosecond laser irradiation exhibited a strong two-photon fluorescence when excited with near-IR femtosecond pulses of the Ti:Sapphire laser. Laser power was kept below 10 mW to avoid generation of additional fluorescent products. The quadratic dependence of the emission intensity on laser power confirms that fluorescence is excited via two-photon absorption (Fig. 2b).

**Figure 2.**
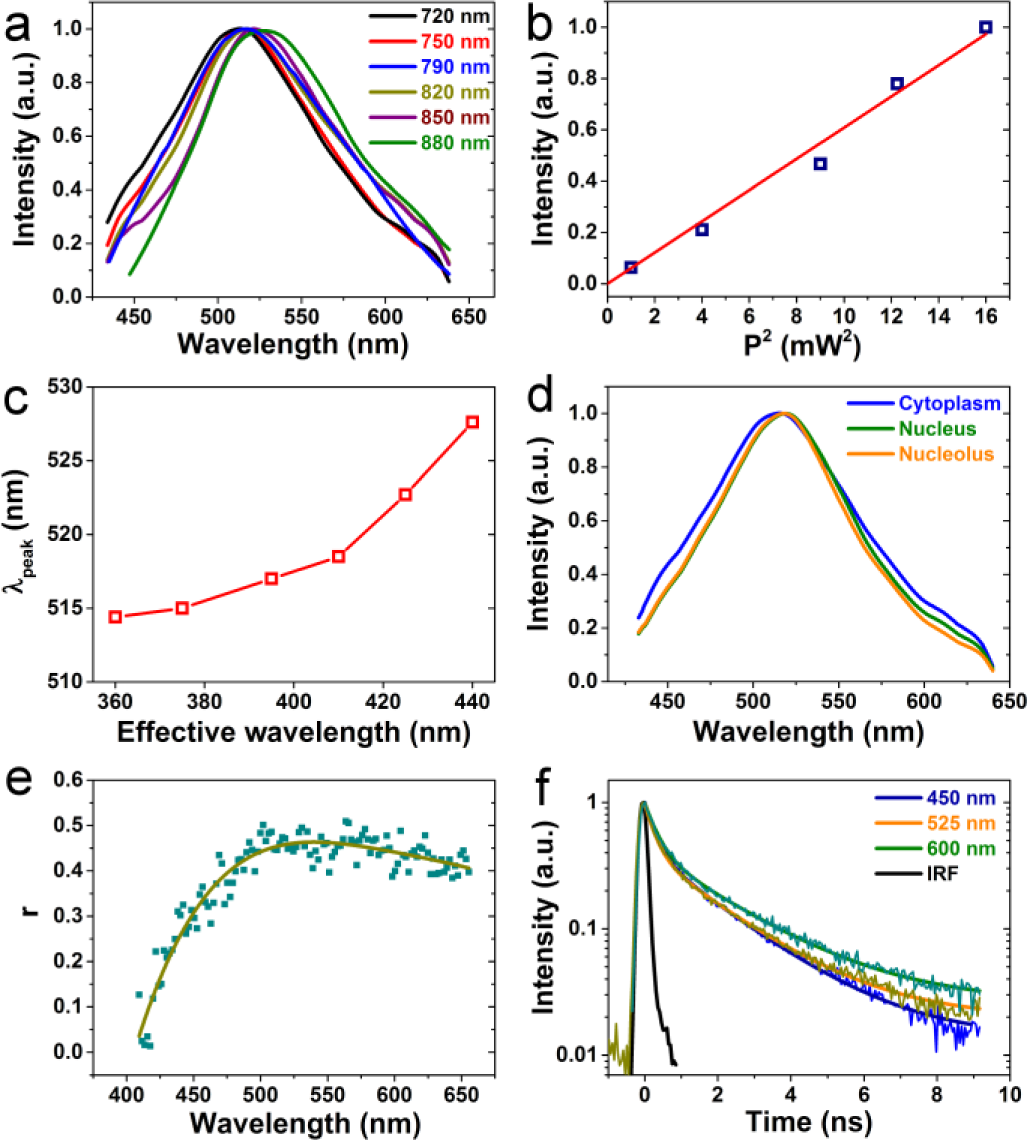
(**a**) Normalized emission spectra of fluorescence from a laser-generated fluorescent spot in the oocyte’s cytoplasm at different excitation wavelengths. Fluorescence was excited by two-photon absorption of femtosecond laser light. (**b**) Emission intensity vs. squared laser power in a fluorescent spot: experimental points and a linear fit (line). (**c**) Emission maximum vs. effective excitation wavelength for a fluorescent spot in the cytoplasm. Effective excitation wavelength is taken equal to a half of the laser wavelength. (**d**) Comparison of normalized emission spectra from laser-generated fluorescent spots in cytoplasm, nucleus and nucleolus of mouse oocyte at 790 nm excitation. (**e**) Fluorescence anisotropy as a function of emission wavelength, two-photon excitation at 790 nm (**f**): Fluorescence decay curves recorded at 450, 525 and 600 nm and their fit with a biexponential decay function, two-photon excitation at 790 nm. A black line represents an instrument response function.

Laser-generated fluorescent material had broad emission spectra with maxima in the green spectral region 500-530 nm (Fig. 2a,d). These spectra overlap with emission of the endogenous cellular fluorophores such as FAD and NADH in blue and green regions, so that laser-generated fluorescence could not be spectrally filtered from cellular autofluorescence. Nevertheless, laser-produced fluorescent spots were well discernible due to the much higher emission intensity. Emission spectra of fluorescent products generated in cytoplasm, nucleus and nucleolus were generally similar, which suggests that the same type of products was generated in all the three regions (Fig. 2d). When the excitation wavelength was tuned from 720 to 880 nm the shape of the emission spectrum remained mostly the same, but its peak wavelength shifted towards longer wavelengths by 15-20 nm. Inset on Figure 2a illustrates monotonous shift of emission maxima with increase of effective excitation wavelength, taken as a half of excitation laser wavelength.

Such behavior indicates inhomogeneous broadening of emission spectra, so we suggest that the fluorescence of laser-generated products consists of contributions from different species or different emission states. This notion is supported by measured fluorescence anisotropy at two-photon excitation that demonstrates a strong dependence on emission wavelength with maximal anisotropy (about 0.45) in the green spectral region near the emission maximum of fluorescent products (Fig. 2e). This manifest wavelength dependence suggests a contribution from several species with different emission anisotropy.

Decay of fluorescence excited by femtosecond pulses in the laser-generated spot was relatively fast with effective fluorescence lifetime smaller than 1 ns (Fig. 1f) which indicates a high non-radiative relaxation rate. Decay curves were non-exponential but could be satisfactorily fitted with a biexponential function having a fast decay time in the range from 0.25 to 0.35 ns and a slow time in the range from 1.9 to 2.4 ns (Table S2). Decay curves depended on emission wavelength; especially with transition to the red edge of the emission spectrum (600 nm) decay became notably slower. In a similar way a red shift of the excitation wavelength from 720 to 880 nm led to slower fluorescence decay (Table S2). In combination with the spectral data these decay kinetics suggest that fluorescence occurs from an ensemble of emissive states different in their emission wavelength and relaxation rate, and states at the red edge of the combined emission spectra possess a longer relaxation time.

An insight into chemical nature of laser-generated products was provided by the spatially resolved Raman spectroscopy. We compared Raman spectra from a non-irradiated cytoplasm material and a relatively large (about 10 μm in diameter) laser-generated fluorescent spot in the cytoplasm (Fig. 3a). The most prominent features in the spectrum of the non-irradiated cytoplasm were broad peaks between 2800 and 3800 cm^−1^ corresponding to stretching vibrations of –CH, −CH2 and −CH3 groups and stretching vibrations of water molecules (Fig. 3b). The same peaks were visible in the Raman spectrum of the laser-generated fluorescent spot, but it also exhibited two strong peaks with maxima at about 1350 and 1590 cm-1 which were not detectable in the normal cytoplasm. These Raman peaks correspond to the known D- and G-bands in graphenic materials. The G band at 1590 cm^−1^ originates from stretching of bonds between sp^2^-hybridized carbon atoms and is commonly observed in graphite, graphene and similar sp^2^ carbon materials. The D band at 1350 cm^−1^ originates from disorder in sp^2^-hybridized carbon, for example edges in carbon sheets or graphite crystals, thus strong D band is typical for graphite crystallites of nanoscale sizes and large surface-to-volume ratio [36]. Both bands are considerably broadened which indicates a high degree of disorder. Finally, a weaker and a broad peak at about 2720 cm^−1^ especially pronounced on the differential spectrum is the second order G’ or 2D band, its position and asymmetric shape are typical for disordered graphite or multi-layer graphene [37]. Background-corrected ratio of D and G band intensities *I(D)/I(G)* is approximately 1. As reported by Tuinistra and Koenig [38] in nanocrystalite graphite this ratio is inversely proportional to the effective crystallite size La: *I*(*D*)/*I*(*G*) = *C*(*λ*)·*La*, where *C(λ)* is a coefficient depending on excitation wavelength, yet this relation is not valid anymore for crystallites smaller than several nanometers nm [36]. Using an empirical expression for *C(λ)* from [36] we arrive to La<20 nm. Thus the Raman spectrum indicates that the laser-generated fluorescent area in the cellular material contains carbon dots-like nanoparticles with nanosized graphitic cores. As a conclusion, the bright fluorescence from laser-generated species is largely attributable to CDs, which are known to be strongly fluorescent, and these CDs are most probably produced from biological molecules of the intracellular material.

**Figure 3.**
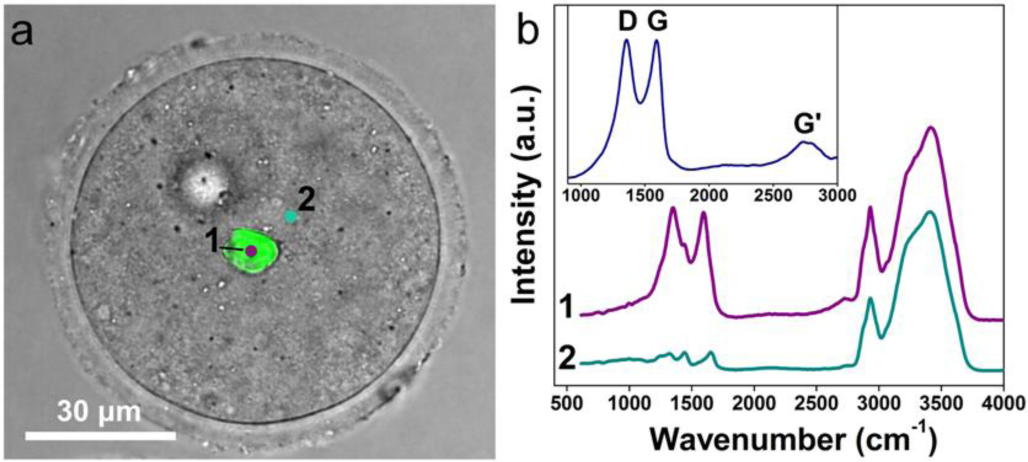
(**a**) An oocyte with a laser-generated fluorescent spot (green area), overlap of brightfield and fluorescent images. (**b**) Normalized Raman spectra from the laser-generated spot (point **1** on (**a**)) and the non-irradiated cytoplasm (point **2** on (**a**)). An inset shows a differential Raman spectrum at the point **1** with subtracted background signal from the cytoplasm.

Formation of carbon dots explains the red shift of emission spectra with increase of the excitation wavelength which is typical for carbon dots fluorescence [39], and is usually attributed to inhomogeneously broadened emission from an ensemble of emissive states with different energies [40]. Also a wavelength-dependent CDs emission decay was reported [41] similar to our observations. A high degree of emission anisotropy suggests that CD fluorescence mostly originates from surface emissive states or dopant atoms [42-43], whereas a smaller anisotropy at the blue edge of the fluorescence spectrum (Fig. 2e) is probably due to a contribution of isotropic emission from the CDs core. The last assumption agrees with an observation that CDs may exhibit luminescence from the carbon core at about 400 nm [44]. Mechanism of laser-induced CDs formation requires a further study, but apparently unlike laser ablation of graphite [45] or other carbon materials it involves a bottom-up synthesis of carbon nanoparticles from organic molecules. A similar bottom-up synthesis of graphene dots from benzene under nanosecond laser irradiation was reported previously [31]. We observed that thermal treatment of the intracellular material using focused CW IR radiation could not reproduce generation of brightly fluorescent species, hence excitation to high-energy states or photoinization and electron plasma generation are likely to play role in carbon dots formation. It is probable that we realize plasma-mediated synthesis [29-30] with a low-density microscale electronic plasma produced by nonlinear photoionization.

Laser generated carbon dots in a cellular material offer a potential for applications in fluorescent imaging as fluorescent “micro-markers”. One of the key problems for bioimaging applications is how long fluorescent spots persist after formation. In order to investigate stability of laser-generated fluorescent spots oocytes after laser irradiation were placed in the CO_2_ incubator for the period up to 24 hours. We observed that larger spots with characteristic diameter of at least several μm remained visible 24 hours after incubation, whereas smaller spots stayed for at least several hours. Figure 4a,b presents images of fluorescent spots generated in different areas of oocytes before and after incubation. In both cases fluorescent spots remained observable after the incubation period and their sizes and brightness remained generally the same. We conclude that laser-synthesized carbon dots stay embedded in the biological matrix in cytoskeleton, adhered to organelles and show no signs of diffuse spreading for at least tens of hours. Their fluorescent properties persist for the same duration which indicates a high degree of chemical stability. Consequently laser-induced generation of fluorescent material can be used to produce fluorescent markers for tracking of intracellular movement and reorganization of cell structure and marking of individual cell in culture.

**Figure 4.**
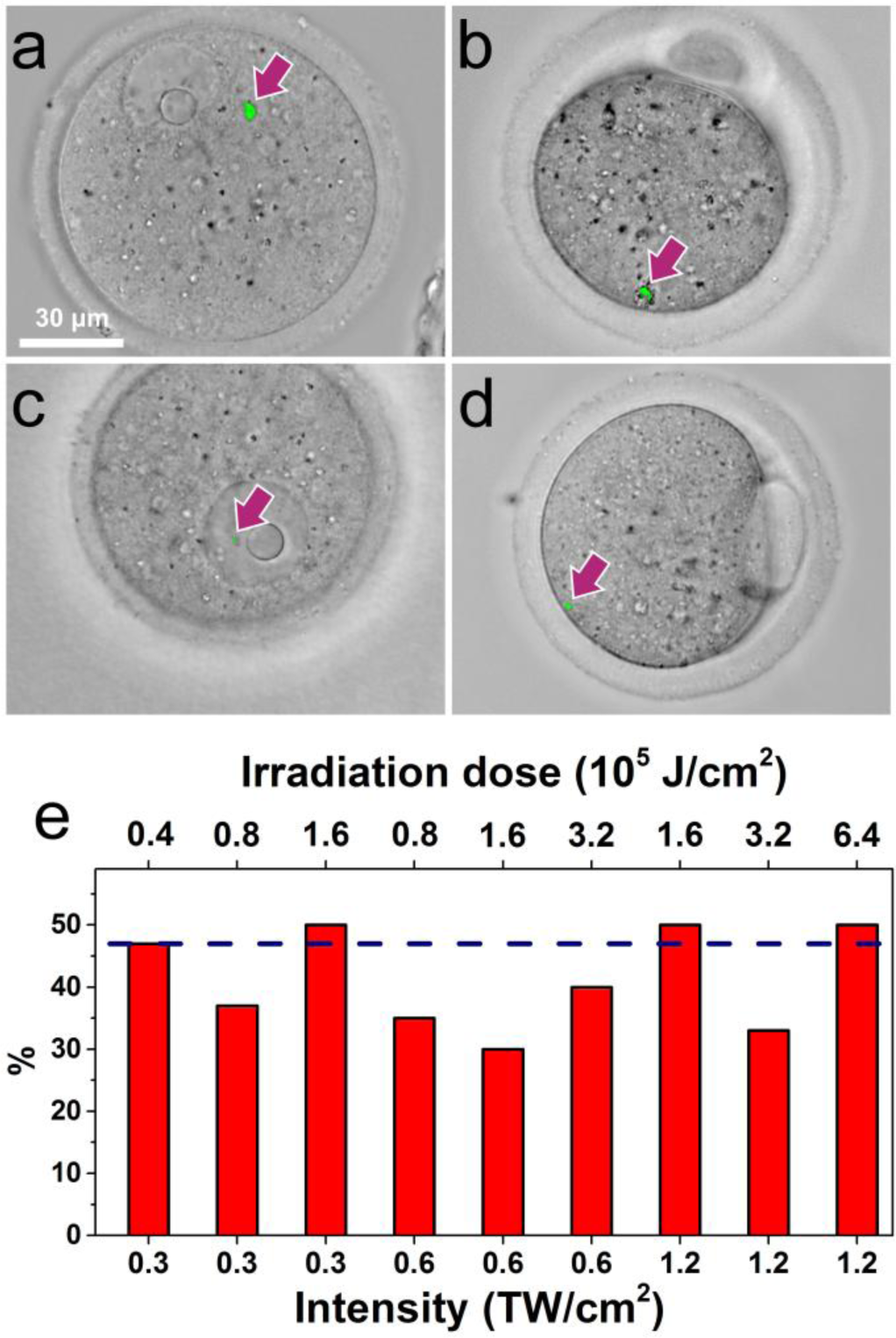
Images of a fluorescent marker in cytoplasm (**a**), (**b**) and nucleus (**c**), (**d**) immediately after laser irradiation (**a**), (**c**) and after a 24 hours incubation (**b**), (**d**). (**e**) Percentage of GV oocytes that maturated to the MII phase after laser exposure of cytoplasm as a function of irradiation parameters. The dashed line shows the maturation percentage in a control group of cells not subjected to a laser exposure.

It is important to note that location of laser-generated spots changed during the incubation period and in case of the spot in Fig.4c the nucleus itself completely dissolved. Still, we were able to track migration of the spot inside the cell using fluorescence images. Thus, Fig.4 provides an example of how laser-generated fluorescent markers can be practically used to visualize reorganization of intracellular material.

Another observation from Figure 4b,d is that the laser-irradiated oocyte maturated from GV to the MII phase, as seen from separation of the polar body. Maturation to MII was also confirmed by observation of the metaphase plate in fluorescent images. Exposure of oocyte material to femtosecond laser pulses produces local heating and also leads to formation of low density electron plasma and production of new chemical species including free radicals and reactive oxygen species [46] which can negatively affect developmental competence of oocytes. Still, these potentially harmful effects did not interfere with oocyte maturation. We quantitatively characterized influence of femtosecond laser irradiation on oocyte development using maturation to the MII phase as a simple criterion of laser-induced damage. Fig. 4e presents the percentage of oocytes that developed to the MII phase during 24 hours after laser irradiation of an arbitrary area in the cytoplasm as a function of irradiation parameters (laser pulse intensity and irradiation dose), detailed statistics are presented in Table S3. Oocytes in the control group were not subjected to laser irradiation and corresponding percentage is shown as a dashed line in Fig. 4e for comparison. The average percentage of oocytes reaching MII in laser-irradiated groups was comparable to the control group. Moreover, increase in pulse energy and exposure time did not have a conspicuous effect on oocyte development. In particular, the percentage of MII oocytes in the groups irradiated with 1.2 TW/cm^2^ laser intensity was roughly the same as in the groups irradiated with 0.3 TW/cm^2^ intensity. As follows from Fig.1e, exposure of cytoplasm to 1.2 TW/cm^2^ pulses generates fluorescent spots with average diameters of 4-5 µm, whereas 0.3 TW/cm^2^ pulses generate almost no fluorescent material. Thus, formation of relatively large fluorescent spots had no discernible influence on ability of GV oocytes to develop to the MII phase. Fluorescent spots generated in this experiment are relatively large compared with typical spots that can be used to track intracellular movement. Consequently, laser generation of fluorescent products within cells induces only a moderate cytotoxic effect and does not affect developmental competence of oocytes, which opens the way for practical applications of this method in live-cell imaging.

## CONCLUSION

In this paper we demonstrated a new technique for fluorescent imaging of living cells which is based on formation of fluorescent products through chemical reactions initiated by nonlinear absorption of femtosecond laser pulses in a miscroscale volume of intracellular material. Fluorescence and Raman spectroscopy indicate that these fluorescent products are at least partly composed of carbon dots with sp^2^ hybridized carbon core. The exact formation mechanism of CDs require a further study, but it is probable that fluorescent carbon dots are produced by low density plasma-mediated carbonization of organic compounds in the cellular material. We have demonstrated that our technique offers an unprecedented control of formation of fluorescent microscale markers within cells: they could be created with high precision in practically any arbitrary point in the intracellular material. The size of the marker can be controlled by tuning the laser power and sizes as small as hundreds of nanometers are achievable, which gives an opportunity to label single organelles. Compared with conventional methods of fluorescent staining our technique does not have an ability to label specific biomolecules in a chemically selective manner, but on the other hand it can spatially selectively target a specific microscale volume within the cell. Laser-generated carbon dots are strongly two-photon active and offer a potential for multiphoton fluorescent imaging. Products of laser-induced chemical reactions do not exhibit noticeable toxicity and the procedure of laser treatment itself does not interfere with cell viability and development. Moreover, laser-produced fluorescent markers exhibit high chemical and structural stability and retain bright fluorescence many hours after formation and thus can be used to track reorganization of cellular material. Laser-induced formation of fluorescent markers within the cell could be used in precision tracking techniques to obtain information about cell cycle, reorganization of the cell structure and intracellular transport.

## ACKNOWLEDGMENTS

This work was supported by the state task for ICP RAS 0082-2018-0005 (registration number AAAA-A18-118020690203-8) and by Russian Foundation for Fundamental research project No. 18-03-01121, and was performed on CKP # 506694 equipment.

## REFERENCES

1. O.S. Wolfbeis. An Overview of Nanoparticles Commonly Used in Fluorescent Bioimaging. Chemical Society Reviews 2015, 44, 4743–4768.

2. U. Resch-Genger, M. Grabolle, S. Cavaliere-Jaricot, R. Nitschke, T. Nann. Quantum Dots Versus Organic Dyes as Fluorescent Labels. Nature Methods 2008, 5, 763.

3. S. Zhu, Q. Meng, L. Wang, J. Zhang, Y. Song, H. Jin, K. Zhang, H. Sun, H. Wang, B. Yang. Highly Photoluminescent Carbon Dots for Multicolor Patterning, Sensors, and Bioimaging. Angewandte Chemie International Edition 2013, 52, 3953–3957.

4. Q.-L. Zhao, Z.-L. Zhang, B.-H. Huang, J. Peng, M. Zhang, D.-W. Pang. Facile Preparation of Low Cytotoxicity Fluorescent Carbon Nanocrystals by Electrooxidation of Graphite. Chemical Communications 2008, 5116–5118.

5. S.-T. Yang, L. Cao, P.G. Luo, F. Lu, X. Wang, H. Wang, M.J. Meziani, Y. Liu, G. Qi, Y.-P. Sun. Carbon Dots for Optical Imaging in Vivo. Journal of the American Chemical Society 2009, 131, 11308–11309.

6. L. Cao, et al. Carbon Dots for Multiphoton Bioimaging. Journal of the American Chemical Society 2007, 129, 11318–11319.

7. X. Xu, R. Ray, Y. Gu, H.J. Ploehn, L. Gearheart, K. Raker, W.A. Scrivens. Electrophoretic Analysis and Purification of Fluorescent Single-Walled Carbon Nanotube Fragments. Journal of the American Chemical Society 2004, 126, 12736–12737.

8. V. Nguyen, L. Yan, J. Si, X. Hou. Femtosecond Laser-Assisted Synthesis of Highly Photoluminescent Carbon Nanodots for Fe3+ Detection with High Sensitivity and Selectivity. Optical Materials Express 2016, 6, 312–320.

9. J. Lu, J.-x. Yang, J. Wang, A. Lim, S. Wang, K.P. Loh. One-Pot Synthesis of Fluorescent Carbon Nanoribbons, Nanoparticles, and Graphene by the Exfoliation of Graphite in Ionic Liquids. ACS Nano 2009, 3, 2367–2375.

10. H. Li, X. He, Y. Liu, H. Yu, Z. Kang, S.-T. Lee. Synthesis of Fluorescent Carbon Nanoparticles Directly from Active Carbon Via a One-Step Ultrasonic Treatment. Materials Research Bulletin 2011, 46, 147–151.

11. C.-B. Ma, et al. A General Solid-State Synthesis of Chemically-Doped Fluorescent Graphene Quantum Dots for Bioimaging and Optoelectronic Applications. Nanoscale 2015, 7, 10162–10169.

12. X. Jia, J. Li, E. Wang. One-Pot Green Synthesis of Optically Ph-Sensitive Carbon Dots with Upconversion Luminescence. Nanoscale 2012, 4, 5572–5575.

13. X. Wang, K. Qu, B. Xu, J. Ren, X. Qu. Microwave Assisted One-Step Green Synthesis of Cell-Permeable Multicolor Photoluminescent Carbon Dots without Surface Passivation Reagents. Journal of Materials Chemistry 2011, 21, 2445–2450.

14. S.Y. Park, H.U. Lee, E.S. Park, S.C. Lee, J.-W. Lee, S.W. Jeong, C.H. Kim, Y.-C. Lee, Y.S. Huh, J. Lee. Photoluminescent Green Carbon Nanodots from Food-Waste-Derived Sources: Large-Scale Synthesis, Properties, and Biomedical Applications. ACS Applied Materials & Interfaces 2014, 6, 3365–3370.

15. I. Canton, G. Battaglia. Endocytosis at the Nanoscale. Chemical Society Reviews 2012, 41, 2718–2739.

16. L. Shang, K. Nienhaus, G.U. Nienhaus. Engineered Nanoparticles Interacting with Cells: Size Matters. Journal of Nanobiotechnology 2014, 12, 5.

17. K. Kettler, K. Veltman, D. van de Meent, A. van Wezel, A.J. Hendriks. Cellular Uptake of Nanoparticles as Determined by Particle Properties, Experimental Conditions, and Cell Type. Environmental Toxicology and Chemistry 2014, 33, 481–492.

18. B. Yameen, W.I. Choi, C. Vilos, A. Swami, J. Shi, O.C. Farokhzad. Insight into Nanoparticle Cellular Uptake and Intracellular Targeting. Journal of Controlled Release 2014, 190, 485–499.

19. P.H. Hemmerich, A.H. von Mikecz. Defining the Subcellular Interface of Nanoparticles by Live-Cell Imaging. PLoS One 2013, 8, e62018.

20. R.R. Gattass, E. Mazur. Femtosecond Laser Micromachining in Transparent Materials. Nature Photonics 2008, 2, 219–225.

21. F. Korte, J. Serbin, J. Koch, A. Egbert, C. Fallnich, A. Ostendorf, B.N. Chichkov. Towards Nanostructuring with Femtosecond Laser Pulses. Applied Physics A 2003, 77, 229–235.

22. M. Malinauskas, A. Žukauskas, S. Hasegawa, Y. Hayasaki, V. Mizeikis, R. Buividas, S. Juodkazis. Ultrafast Laser Processing of Materials: From Science to Industry. Light: Science & Applications 2016, 5, e16133.

23. B.H. Cumpston, et al. Two-Photon Polymerization Initiators for Three-Dimensional Optical Data Storage and Microfabrication. Nature 1999, 398, 51.

24. M.T. Raimondi, S.M. Eaton, M.M. Nava, M. Laganà, G. Cerullo, R. Osellame. Two-Photon Laser Polymerization: From Fundamentals to Biomedical Application in Tissue Engineering and Regenerative Medicine. Journal of Applied Biomaterials & Functional Materials 2012, 10, 56–66.

25. T. Tanaka, A. Ishikawa, S. Kawata. Two-Photon-Induced Reduction of Metal Ions for Fabricating Three-Dimensional Electrically Conductive Metallic Microstructure. Applied Physics Letters 2006, 88, 081107.

26. S. Maruo, T. Saeki. Femtosecond Laser Direct Writing of Metallic Microstructures by Photoreduction of Silver Nitrate in a Polymer Matrix. Optics Express 2008, 16, 1174–1179.

27. A.V.a.V. Venugopalan. Mechanisms of Pulsed Laser Ablation of Biological Tissues. Chem. Rev. 2003, 103, 577–644.

28. A. Vogel, J. Noack, G. Hüttman, G. Paltauf. Mechanisms of Femtosecond Laser Nanosurgery of Cells and Tissues. Applied Physics B 2005, 81, 1015–1047.

29. J. Wang, C.-F. Wang, S. Chen. Amphiphilic Egg-Derived Carbon Dots: Rapid Plasma Fabrication, Pyrolysis Process, and Multicolor Printing Patterns. Angewandte Chemie International Edition 2012, 51, 9297–9301.

30. X. Huang, Y. Li, X. Zhong, A.E. Rider, K. Ostrikov. Fast Microplasma Synthesis of Blue Luminescent Carbon Quantum Dots at Ambient Conditions. Plasma Processes and Polymers 2015, 12, 59–65.

31. K. Habiba, V.I. Makarov, J. Avalos, M.J.F. Guinel, B.R. Weiner, G. Morell. Luminescent Graphene Quantum Dots Fabricated by Pulsed Laser Synthesis. Carbon 2013, 64, 341–350.

32. R. Galli, O. Uckermann, E.F. Andresen, K.D. Geiger, E. Koch, G. Schackert, G. Steiner, M. Kirsch. Intrinsic Indicator of Photodamage During Label-Free Multiphoton Microscopy of Cells and Tissues. PLoS One 2014, 9, e110295.

33. Q. Sun, Z. Qin, W. Wu, Y. Lin, C. Chen, S. He, X. Li, Z. Wu, Y. Luo, J.Y. Qu. In Vivo Imaging-Guided Microsurgery Based on Femtosecond Laser Produced New Fluorescent Compounds in Biological Tissues. Biomedical Optics Express 2018, 9, 581–590.

34. Z. Qin, Q. Sun, Y. Lin, S. He, X. Li, C. Chen, W. Wu, Y. Luo, J.Y. Qu. New Fluorescent Compounds Produced by Femtosecond Laser Surgery in Biological Tissues: The Mechanisms. Biomedical Optics Express 2018, 9, 3373–3390.

35. A.A. Astaf’ev, A.D. Zalesskii, A.M. Shakhov, A.A. Osychenko, V.A. Nadtochenko. Formation of Light-Absorbing Centers Induced in Cytoplasm of Mouse Embryos by Femtosecond Pulsed near-Infrared Radiation. High Energy Chemistry 2016, 50, 421–423.

36. M.S. Dresselhaus, A. Jorio, A.G. Souza Filho, R. Saito. Defect Characterization in Graphene and Carbon Nanotubes Using Raman Spectroscopy. Philosophical Transactions of the Royal Society A: Mathematical, Physical and Engineering Sciences 2010, 368, 5355–5377.

37. L.M. Malard, M.A. Pimenta, G. Dresselhaus, M.S. Dresselhaus. Raman Spectroscopy in Graphene. Physics Reports 2009, 473, 51–87.

38. F. Tuinstra, J.L. Koenig. Raman Spectrum of Graphite. The Journal of Chemical Physics 1970, 53, 1126–1130.

39. Y.-P. Sun, et al. Quantum-Sized Carbon Dots for Bright and Colorful Photoluminescence. Journal of the American Chemical Society 2006, 128, 7756–7757.

40. G.E. LeCroy, F. Messina, A. Sciortino, C.E. Bunker, P. Wang, K.A.S. Fernando, Y.-P. Sun. Characteristic Excitation Wavelength Dependence of Fluorescence Emissions in Carbon “Quantum” Dots. The Journal of Physical Chemistry C 2017, 121, 28180–28186.

41. S. Khan, A. Gupta, N.C. Verma, C.K. Nandi. Time-Resolved Emission Reveals Ensemble of Emissive States as the Origin of Multicolor Fluorescence in Carbon Dots. Nano Letters 2015, 15, 8300–8305.

42. M.O. Dekaliuk, O. Viagin, Y.V. Malyukin, A.P. Demchenko. Fluorescent Carbon Nanomaterials: “Quantum Dots” or Nanoclusters? Physical Chemistry Chemical Physics 2014, 16, 16075–16084.

43. P. Jing, D. Han, D. Li, D. Zhou, L. Zhang, H. Zhang, D. Shen, S. Qu. Origin of Anisotropic Photoluminescence in Heteroatom-Doped Carbon Nanodots. Advanced Optical Materials 2017, 5, 1601049.

44. N. Dhenadhayalan, K.-C. Lin, R. Suresh, P. Ramamurthy. Unravelling the Multiple Emissive States in Citric-Acid-Derived Carbon Dots. The Journal of Physical Chemistry C 2016, 120, 1252–1261.

45. S.-L. Hu, K.-Y. Niu, J. Sun, J. Yang, N.-Q. Zhao, X.-W. Du. One-Step Synthesis of Fluorescent Carbon Nanoparticles by Laser Irradiation. Journal of Materials Chemistry 2009, 19, 484–488.

46. U.K. Tirlapur, K. König, C. Peuckert, R. Krieg, K.-J. Halbhuber. Femtosecond near-Infrared Laser Pulses Elicit Generation of Reactive Oxygen Species in Mammalian Cells Leading to Apoptosis-Like Death. Experimental Cell Research 2001, 263, 88–97.

