## Supplementary information for "Live cell bioimaging with carbon dots produced in situ by femtosecond laser from intracellular material"

### Supporting information

#### Material and methods

*Laser setup.* Femtosecond laser pulses with a 80 MHz repetition rate and energy up to 25 nJ were generated by a Titanium-Sapphire oscillator (Tsunami, Spectra-Physics) pumped by a DPSS Nd:$YVO_4$ CW laser with 532 nm wavelength (Millennia Prime 6sJ, Spectra-Physics). The central wavelength varied from 690 to 930 nm, typically the wavelength of 790 nm was used. The average laser power was tuned with a polarizing attenuator consisting of a half-wave plate and a polarizing cube. The maximum average power before the objective lens was 700 mW. The laser pulse trains were coupled to an inverted optical microscope Olympus IX71 by Thorlabs FESH0750 dielectric filter mounted at $45^0$ and then focused by a 40x 0.75NA objective lens (UPlanFLN, Olympus) on a sample, which was placed on a 3-axis stage. Laser radiation was focused on the sample with a 40x 0.75NA lens (UPlanFLN, Olympus). The laser beam completely filled the aperture of the objective. Taking the beam quality factor $M^2$ as 1 the focal spot diameter is estimated as $d = 1.22\lambda/NA$ ~ 1.3 μm and confocal parameter as $b=\pi d^2/2\lambda = 3.28$ μm. The pulse duration in the focal plane was measured by the Avesta AA-M autocorrelator and was about 25 fs. The SF11 prism compressor was used to compensate for the group velocity dispersion in the objective lens and other optical elements. The length of the pulse trains was determined by opening time of a mechanical shutter (SH05, Thorlabs) unblocking the femtosecond laser beam for a time up to 200 ms. The brightfield image of the sample was recorded by a XIMEA xiQ MQ013MG-ON CMOS camera or by XIMEA xiD MD061CU-SY CCD camera, mounted on the microscope.

*Fluorescence and Raman registration*. We used two methods to record fluorescence signal from the sample. In the first method fluorescence was excited by single-photon absorption of laser diode radiation with λ = 462 nm (Nichia NDB7675), which was coupled to the Olympus microscope and focused by the same objective lens as femtosecond radiation. The laser diode beam was focused on the objective lens entrance pupil to give an about 500 μm wide and uniform field of illumination. The fluorescent image was also recorded by the same cameras as described above with exposure time up to 1 second. The excitation light was cut off by the long-pass dielectric filter Thorlabs FELH500.

In the second method, two-photon fluorescence was excited by two-photon absorption of femtosecond laser pulses focused by the objective lens. In order to avoid damage to the biological material or chemical reactions induced by laser radiation its average power was reduced below 10 mW. Fluorescence signal was collected by the same objective lens and directed to the microscope's side port via a beamsplitter cube and then was coupled to an Acton SP300i monochromator and then to a PI-MAX 2 CCD camera (Princeton Instruments) used to record the fluorescence spectra or to a PMT of the time-correlated single photon counting system SPC-730 (Becker & Hickl GmbH) for detecting the fluorescence decay kinetics. For recording spatially averaged spectra and decay kinetics the sample was raster scanned relative to the focused laser beam using a piezoelectric scanning stage (NT-MDT); the scan area was typically several μm wide. A linear polarizer could be additionally installed between the microscope and the monochromator to record emission anisotropy.

Raman scattering of the oocyte material was excited with a 532-nm DPSS CW laser (Coherent). Laser power at the focal spot was kept between 1 and 10 mW to avoid laser damage to the sample. Raman signal was collected by the objective and registered by the Renishaw 1000B micro-Raman spectrometer attached to the microscope.

*Oocyte collection*. CBA/C57Bl female hybrid mice aged 1-1.5 month were injected 10 IU pregnant mare serum gonadotropin (A036A02 "Intervet") 48 hours before oocyte collection. Injected females were killed by cervical dislocation. The ovaries were recovered from mice and placed into 2 ml warm PBS medium (D4031 “Sigma”) in the 35 mm Petri dishes (353001 “Falcon”). Cumulus-oocyte complexes (COCs) were extracted from ovaries and placed in M2 medium (M7167 “Sigma”) containing 0.1% hyaluronidase (H4272 “Sigma”) to remove cumulus cells. Then oocytes were washed in PBS medium and moved about into 50 μl drop of PBS medium at Petri dish with glass and center hole (100350 “SPL Lifesciences”), covered with 2.5-3 ml mineral oil (M8410 “Sigma”) and immediately used for experiments.

After laser treatment oocytes were washed in M2 medium, transferred to four-well plastic dishes (30004 “SPL Lifesciences”) with 0.7 ml IVM medium and cultivated *in vitro* in $CO_2$-incubator at 37 $^{o}$C with 5% $CO_2$. IVM culture medium composed of DMEM (C420 “PanEco”) supplemented with 15% fetal bovine serum (I31966-021 “Gibco”), 1.5 IU/ml gentamycin (G1272 “Sigma”) and 1 IU/ml pregnant mare serum gonadotropin (A036A02 “Intervet”). After overnight cultivation oocytes were examined for development to the metaphase II stage detected by the presence of a polar body and metaphase plate under light microscopy.

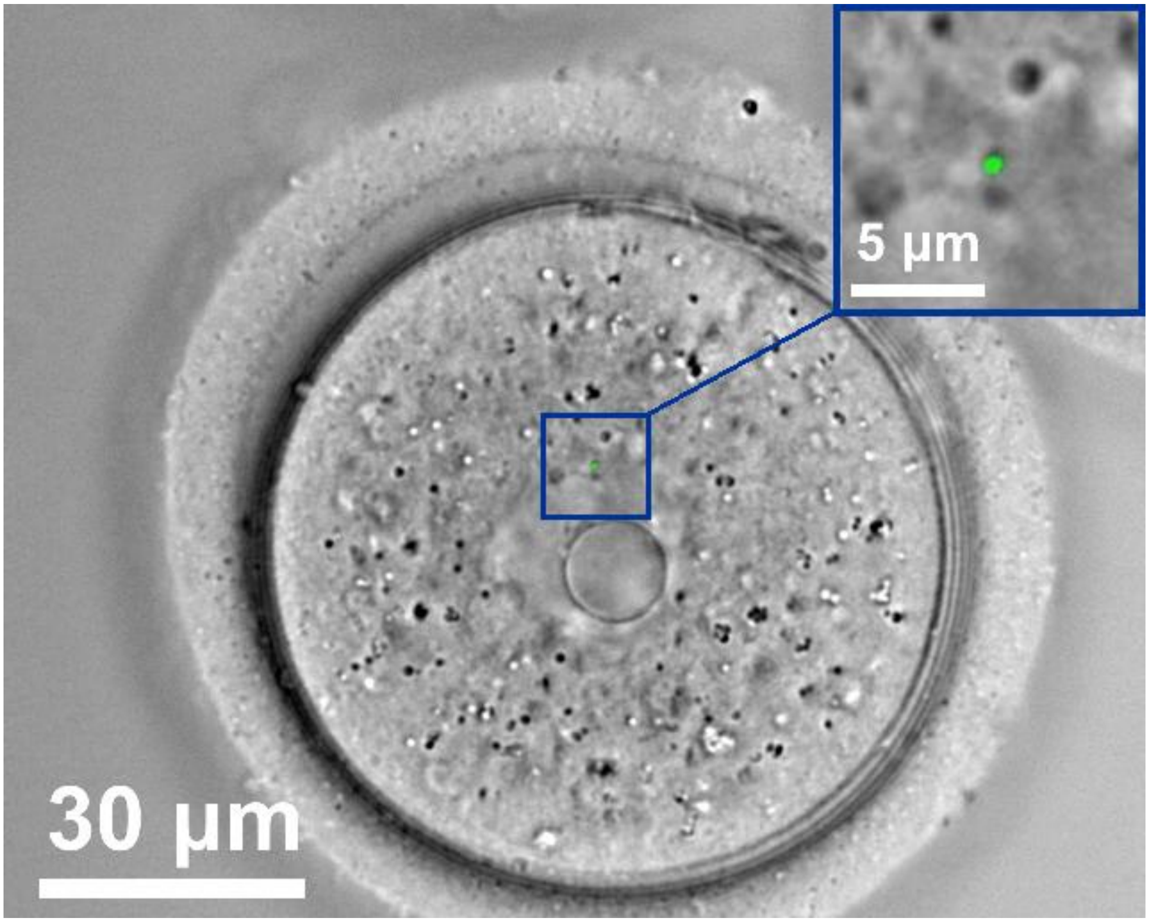


**Figure S1**. A submicron fluorescent spot formed in the oocyte cytoplasm after an irradiation with a femtosecond pulse train. Spot diameter is 0.8 µm.

| Area | Excitation | Emission | $A_1$ | $T_1$, ns | $A_2$ | $T_2$, ns |
|---|---|---|---|---|---|---|
| Non-irradiated | 790 | 525 | 0.46 | 0.43 | 0.54 | 2.21 |
| Non-irradiated | 705 | 525 | 0.54 | 0.44 | 0.46 | 2.07 |
| Irradiated | 790 | 525 | 0.59 | 0.28 | 0.41 | 1.93 |
| Irradiated | 790 | 450 | 0.60 | 0.27 | 0.40 | 1.97 |
| Irradiated | 790 | 600 | 0.59 | 0.33 | 0.41 | 2.20 |
| Irradiated | 705 | 525 | 0.57 | 0.26 | 0.43 | 2.07 |
| Irradiated | 880 | 525 | 0.53 | 0.31 | 0.47 | 2.4 |

**Table S2**. Parameters of a double-exponential fit of measured decay curves in irradiated and non-irradiated areas of the cytoplasm at different excitation and emission wavelengths.

| Irradiation parameters | Number of oocytes in the group | Number of oocytes that developed to MII phase | % of maturation |
|---|---|---|---|
| 0.3 $TW/cm^2$, 40 $kJ/cm^2$ | 30 | 14 | 47 |
| 0.3 $TW/cm^2$, 80 $kJ/cm^2$ | 30 | 11 | 37 |
| 0.3 $TW/cm^2$, 160 $kJ/cm^2$ | 30 | 15 | 50 |
| 0.6 $TW/cm^2$, 80 $kJ/cm^2$ | 20 | 7 | 35 |
| 0.6 $TW/cm^2$, 160 $kJ/cm^2$ | 30 | 9 | 30 |
| 0.6 $TW/cm^2$, 320 $kJ/cm^2$ | 40 | 16 | 40 |
| 1.2 $TW/cm^2$, 160 $kJ/cm^2$ | 30 | 15 | 50 |
| 1.2 $TW/cm^2$, 320 $kJ/cm^2$ | 30 | 10 | 33 |
| 1.2 $TW/cm^2$, 640 $kJ/cm^2$ | 20 | 10 | 50 |
| Control | 230 | 108 | 47 |

**Table S3**. Statistics on GV oocytes maturation to the MII phase after femtosecond laser irradiation of arbitrarily chosen area of the cytoplasm as a function of irradiation parameters (laser intensity and irradiation dose). Oocytes in the control group were not subjected to laser irradiation but otherwise were kept under the same conditions.
